# An exploratory analysis of 4844 withdrawn articles and their retraction notes

**DOI:** 10.1101/2021.09.30.462625

**Authors:** Catalin Toma, Liliana Padureanu

## Abstract

The objective of our study was to obtain an updated image of the dynamic of retractions and retraction notes, retraction reasons for questionable research and publication practices, countries producing retracted articles, and the scientific impact of retractions by studying 4844 PubMed indexed retracted articles published between 2009 and 2020 and their retraction notes.

**RESULTS:** Mistakes/inconsistent data account for 32% of total retractions, followed by images(22,5%), plagiarism(13,7%) and overlap(11,5%).

Thirty countries account for 94,79% of 4844 retractions. Top five are: China(32,78%), United States(18,84%), India(7,25%), Japan(4,37%) and Italy(3,75%).

The total citations number for all articles is 140810(Google Scholar), 96000(Dimensions).

Average exposure time(ET) is 28,89 months. Largest ET is for image retractions(49,3 months), lowest ET is for editorial errors(11,2 months).

The impact of retracted research is higher for Spain, Sweden, United Kingdom, United States, and other nine countries and lower for Pakistan, Turkey, Malaysia, and other six countries, including China.

**CONCLUSIONS:** Mistakes and data inconsistencies represent the main retraction reason; images and ethical issues show a growing trend, while plagiarism and overlap still represent a significant problem. There is a steady increase in QRP and QPP article withdrawals. Retraction of articles seems to be a technology-dependent process.

The number of citations of retracted articles shows a high impact of papers published by authors from certain countries. The number of retracted articles per country does not always accurately reflect the scientific impact of QRP/QPP articles.

The country distribution of retraction reasons shows structural problems in the organization and quality control of scientific research, which have different images depending on geographical location, economic development, and cultural model.

## Introduction

*“We publish this Note at this time because in the years that have passed since our original publication, and even now, the 1959 note has been quoted as a worthy piece of evidence. We ask to be spared the further embarrassment of having that earlier work cited in the reputable literature, and we hope we can spare other authors the labors of attempting to rationalize our aberrant data.”* [1]

The above withdrawal note reflects an issue that continues to be relevant: a scientific article, once published, continues to exert influence and have an impact on the scientific community even if it is proven, after publication, that it contains errors that raise doubts on the conclusions, invalidating them totally or partially.

In 1756 appeared what is most likely the first documented withdrawal of scientific work [2], followed, over 150 years later, by that of Whelden [3] and over time, by many others:

- unreproducible research of questionable scientific value persisted for decades before disappearing from the literature, although it was demonstrated to be wrong shortly after its publication [4] ;
- spectacular results were based on either a laboratory error or an attempt by a researcher to report positive preliminary results [5];
- discoveries presented in an article with over 1000 citations in Google Scholar were questioned one month after publication, following an inspection that found severe problems of research methodology and that today would lead to a very rapid rejection or even would make the article impossible to publish [6];
- for the first time, in addition to the retraction of published papers, the perpetrator of scientific fraud (the Poehlman case) was sentenced to prison [7];
- hundreds of articles were withdrawn due to data fabrication, Schon [8], Hwang [9], Boldt, Reuben, Fuji [10] being just a few examples.

Research misconduct/questionable research practices(QRP) in various forms (from 1,97% up to 72%) within a scientific community has been reported by several studies [11–14].

The costs generated by QRP can reach considerable amounts, conservatively evaluated, in 2010, to cost over 100 million USD only for the cases investigated by O.R.I. in the United States [15] or 400000 USD / article in 2014 [16].

Indirect costs can also increase if no action is taken, such as the funding of research in which contaminated cell lines are being used [17].

Apart from the fairly specific situations in which the scientific value of some articles is questioned, there is an area of questionable publication practices(QPP) in which the reasons for rejection are based instead on non-compliance with ethical standards(ethical writing) and legal regulations (copyright issues): plagiarism, overlap, authorship [18–21]. While plagiarism is rejected and considered a form of scientific misconduct, text re-use/recycling/self-plagiarism is still under debate about the quantity and type of recycled materials[21], and the decision to retract a scientific paper is mainly an editorial one.

If the mechanisms that should prevent the generation, perpetuation, and dissemination of QRP/QPP in biomedical research do not work correctly, there are situations in which harm can be caused to patients, the scientific community, research institutions, funding bodies, publishers, and scientific journals in which the results are published [22–24].

Numerous articles have addressed the subject of retractions in recent years. Similar to papers that used other databases, those that used PubMed / Medline as their sole source of data[25–38] have shown an increasing trend in the number of retracted papers, a diversification of motives and an increased interest of journals and publishers in correcting the scientific literature.

The articles withdrawn from the scientific literature indexed in PubMed, although in a small proportion to the total volume indexed in this database (0.04% for the period 01.01.2009-31.12.2020), are a problem not only by question marks which they can raise on the integrity of the scientific research as a whole but also by the impact on the scientific community, which can use or invest in ideas, methods or data invalidated a few years after publication.

Our study aims to perform an exploratory analysis of human health-related papers withdrawn from the literature indexed in PubMed / Medline and published in the period 2009-2020 focused mainly on:

- dynamic of retracted articles and retraction notes for the period 2009-2020
- retraction reasons
- countries producing QRP/QPP articles
- the scientific impact of retracted papers;

## Materials and methods

### Information sources

- PubMed – PubMed (nih.gov)
- Google Scholar – citations https://scholar.google.com
- Dimensions – https://dimensions.ai
- Scopus – https://scopus.com

Several elements determined the selection of Pubmed as the unique source of information:

- focus on biomedical journals
- unrestricted access
- the existence of a dedicated keyword for withdrawn articles, Retracted Publication [P.T.];
- the link between the withdrawn article and the withdrawal note, Retraction of Publication [P.T.];
- the possibility of exporting the data in csv format in our own database for the individual analysis of each article and the corresponding retraction note.

Google Scholar was used due to its free nature and the best coverage of citations [39].

Dimensions database was used due to free access and provision, in addition to the total number of citations of the number of citations from the last two years (compared to the current date).

SCOPUS (Elsevier) was used to obtain information about journal publishers and the journal impact factor(CiteScore).

### Articles retrieval and extracted information

Withdrawn articles were identified in PubMed using “Retracted Publication [PT]” search without date restrictions. The data were downloaded in csv format and imported into an application developed for analysis by the author.

Data analysis period: 20.07.2020 – 31.05.2021

Last date of data import from PubMed: 30.01.2021

The period analyzed: 01.01.2009 - 31.12.2020 (taking into account the year of publication as recorded in PubMed).

The processing date was noted for each item.

#### Inclusion criteria

the field of study or the subject studied is related or may have an impact on human health (mentioned in the text of the article)

#### Exclusion criteria

the field of study or the studied subject is not related to human health (chemistry, agriculture, veterinary medicine, industrial products, ecology without mentioning in the article some implications with human health) proceedings volumes with no specific retractions mentioned, clinical practice guidelines withdrawn for updates or unspecified reasons, misclassified retraction notes.

Each item and withdrawal note were analyzed, the extracted data is grouped into four sections.

##### Authors and countries

**Table 1.**
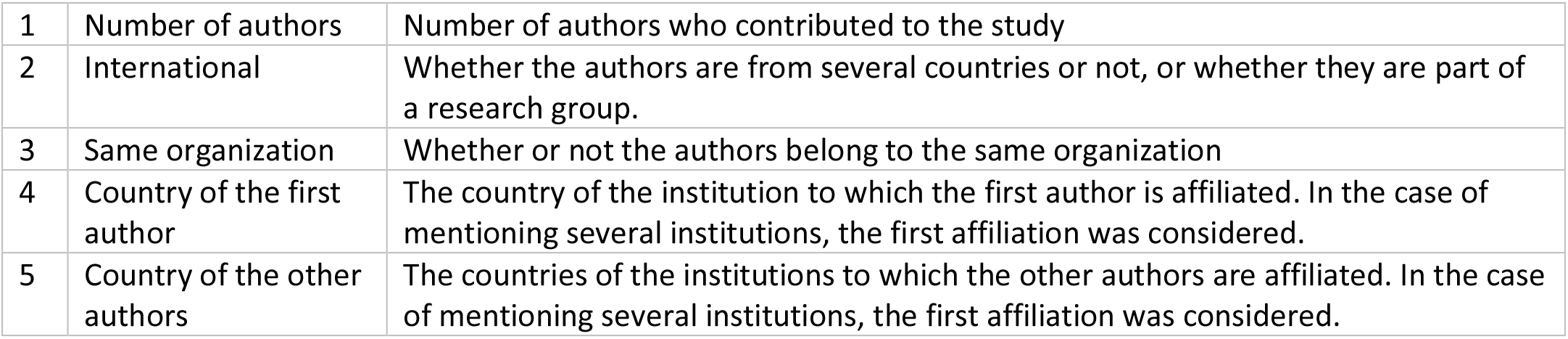
Author and country-related variables.

|  |  |  |
| --- | --- | --- |
| 1 | Number of authors | Number of authors who contributed to the study |
| 2 | International | Whether the authors are from several countries or not, or whether they are part of a research group. |
| 3 | Same organization | Whether or not the authors belong to the same organization |
| 4 | Country of the first author | The country of the institution to which the first author is affiliated. In the case of mentioning several institutions, the first affiliation was considered. |
| 5 | Country of the other authors | The countries of the institutions to which the other authors are affiliated. In the case of mentioning several institutions, the first affiliation was considered. |

##### Retraction details

**Table 2.** Retraction-related variables. (RN = Retraction Note; IRB = Institutional Review Board; ORI = Office of Research Integrity)

|  |  |  |
| --- | --- | --- |
| 6 | Exposure time | the time, measured in months, between the date of withdrawal of the article and the date of publication (latest data) |
| 7 | PMID of retraction note | The PubMed identifier of the withdrawal |
| 8 | Date of RN | Date of retraction note |
| 9 | Doi of RN | The DOI identifier of the withdrawal note |
| 10 | Paper type | Paper type(journal article, proceedings, book chapter) |
| 11 | Study type | case reports, letters/editorials, research article, review, randomized trial, meta-analysis, methods/methodology |
| 12 | Clinical | Source of data used in the article (human subjects, tissues, laboratory, big data) |
| 13 | Available online | If the entire article is online / not / paywalled |
| 14 | Publisher | The journal publisher as reported in Scopus. In case of publisher change (by a takeover of the journal), the publisher was considered from the moment of carrying out this study. |
| 15 | Retract reason | Main retraction reasons: authorship, editorial, ethics, fraud, images, mistakes and or inconsistent data, overlap, plagiarism, property or legal concerns, other, unclear |
| 16 | Retraction details | Text of retraction note (up to 2000 characters) |
| 17 | Image reasons | Details for image related retractions (copyright, duplication, ethical, fabrication, manipulation, overlap, plagiarized, unreliable, wrong) |
| 18 | Reason subcategory | Second level reasons, used when information is available in the retraction note: conflict of interest, duplicate, fabricated data, falsified information, fraudulent peer review, no raw data, no IRB approval, no patient consent, publication w/o approval, reproducibility, research misconduct, research outsourcing, other. |
| 19 | Requested by/Involving | Who asked for/was involved in article retraction.<br>Authors, publisher, editorial (editors or editorial board), institution, O.R.I., other, not specified |

##### Collateral damage

**Table 3.** Collateral damage variables.

|  |  |  |
| --- | --- | --- |
| 20 | R.N. info | Whether or not the withdrawal note contains information related to the individual involvement of the authors |
| 21 | 1st author | The first author involved |
| 22 | Corresponding author | The corresponding author involved |
| 23 | Authors involved | How many authors were involved |
| 24 | Innocent casualties | How many innocent? |

##### Citations

**Table 4.**
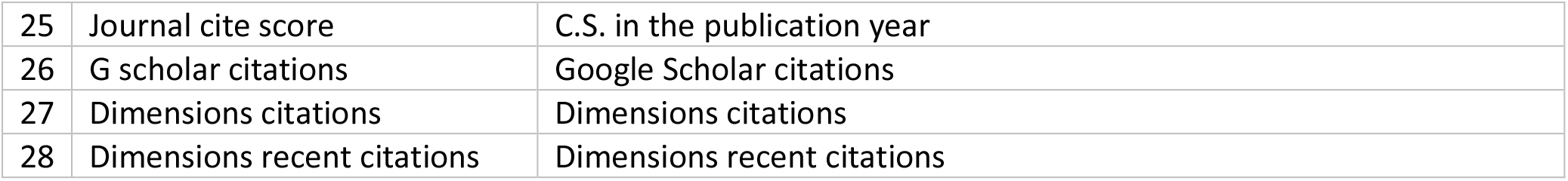
Citations-related variables.

### Definitions

In order to define a flexible taxonomy for the main retraction reasons categories and subcategories, we have considered a series of previously published articles [31,38,40–42].

#### Retraction reasons

**Table 5.** Main retraction reasons.

|  |  |
| --- | --- |
| AUTHORSHIP | - disputes/disagreements between authors related to the content of the article<br>- publication of the article without the consent or knowledge of all authors<br>- listing as authors of people who are not aware of the research<br>- changing authors after article submission |
| EDITORIAL | - duplicate publication of an article in the same journal<br>- other editorial errors (publication before peer review, publication of a wrong version) |
| ETHICS | - lack of patient / patients' consent<br>- withdrawal of consent by the patient/patients after the publication of the article<br>- lack of I.R.B. approval from the institution where the research was conducted/non-compliance with the protocol<br>- failure to comply with the protocol for which the initial I.R.B. approval was obtained<br>- violation of other ethical norms(conflicts of interest, laboratory animals ethics)<br>- publication of an article in which data are used without approval(non-ethical use of data) |
| FRAUD | - manipulating the peer review process by different methods (fake email addresses, fake peer review reports)<br>- forged information (authors, grants, forged ethics agreements, forged consent)<br>- the use in the article of unauthorized or illegally obtained information |
| IMAGES | - any situation in which one of the reasons for withdrawing the article was related to the images used in it |
| MISTAKES/INCONSISTENT DATA | - any type of error that invalidates the results in the article<br>- fabrication of data or results |
| OVERLAP | - re-use of texts, data, or other elements of the article contained in an article published by the same author or group of authors (does not apply to images, which are analyzed separately)<br>- publishing the same article in two or more different journals |
| PLAGIARISM | - use of any information contained in an article already published by other authors (does not apply to images, which are analyzed separately) |
| PROPERTY OR LEGAL CONCERNS | - article in which information protected by copyright or other legal means is used may entail legal liability for the author/publisher/publisher |
| OTHER | - other reasons that do not fall into those described above |

#### Secondary reasons

**Table 6.** Secondary retraction reasons.

|  |  |
| --- | --- |
| CONFLICT OF INTEREST | - conflicts of interest not declared |
| DUPLICATE | - Duplicate publication(editorial mistake) or duplicate submission(author mistake) |
| FABRICATED DATA | - Subcategory of main category Mistakes/Inconsistent data for articles retracted for data fabrication |
| FALSIFIED INFORMATION | - False affiliations, false contact details, falsified documents, credentials, others |
| FRAUDULENT PEER REVIEW | - circumvention or hacking of peer review process |
| NO DATA PROVIDED | - no raw data for the study were available or made available by the authors<br>- no raw data were found following the institutional investigation |
| NO IRB APPROVAL | - nonexistent I.R.B. approval for the study<br>- Study not respecting the protocol approved by IRB |
| NO PATIENT CONSENT | - patient consent incomplete, not given, or withdrawn |
| PUBLICATION W/O APPROVAL | - no publication approval from other authors(coauthor, senior author), institution, funder, or company |
| REPRODUCIBILITY | - reporting by the authors of the impossibility to replicate, totally or partially, the results presented in the article<br>- reporting, by other authors, of the impossibility to replicate the research results from the article |
| RESEARCH MISCONDUCT | - research misconduct was reported by one of the institutions where the research was conducted, by one of the coauthors, or by a 3rd party |
| RESEARCH OUTSOURCING/TEMPLATED | - research performed partially/totally by a 3rd party<br>- templates being used for research reporting |
| OTHER | - various reasons which do not fit within an existing subcategory |

#### Who was involved/requested the retraction

**Table 7.**
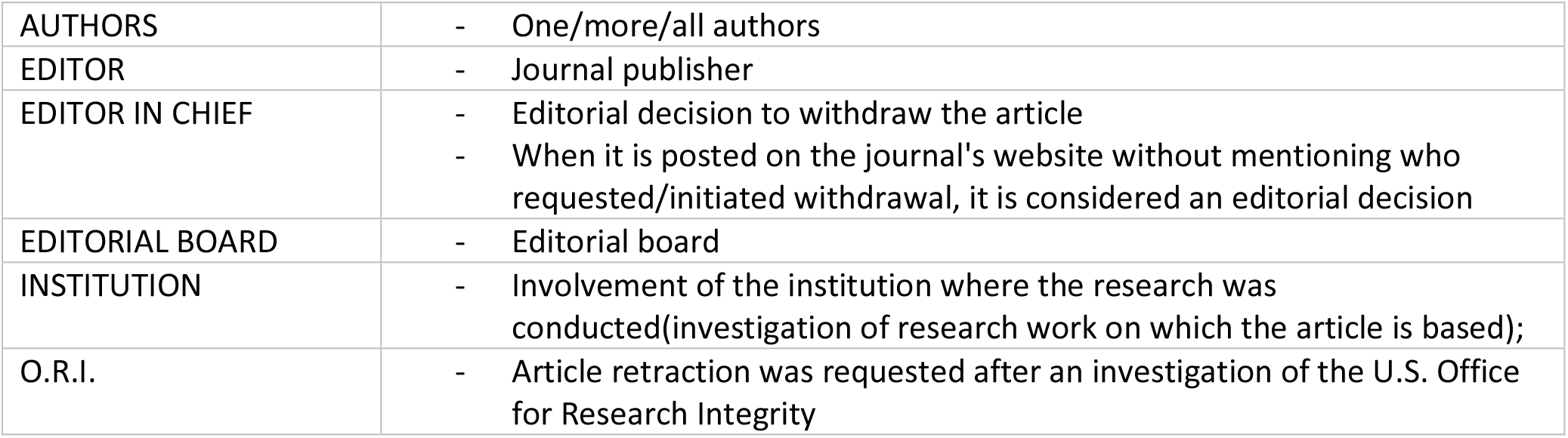
Who was involved in the retraction process.

|  |  |
| --- | --- |
| AUTHORS | - One/more/all authors |
| EDITOR | - Journal publisher |
| EDITOR IN CHIEF | - Editorial decision to withdraw the article<br>- When it is posted on the journal's website without mentioning who requested/initiated withdrawal, it is considered an editorial decision |
| EDITORIAL BOARD | - Editorial board |
| INSTITUTION | - Involvement of the institution where the research was conducted(investigation of research work on which the article is based); |
| O.R.I. | - Article retraction was requested after an investigation of the U.S. Office for Research Integrity |

The data were exported and analyzed in IBM SPSS (IBM SPSS Statistics for Windows 2018).

## Results

### Retracted articles and retraction notes

A total of 5619 retracted papers were retrieved by a PubMed search for the period 2009-2020. Of these, 775 were excluded and 4844 analyzed. The distribution for the period 2009-2020 of the withdrawn articles and the withdrawal notes is presented in Figure 1 and table 8.

**Figure 1.**
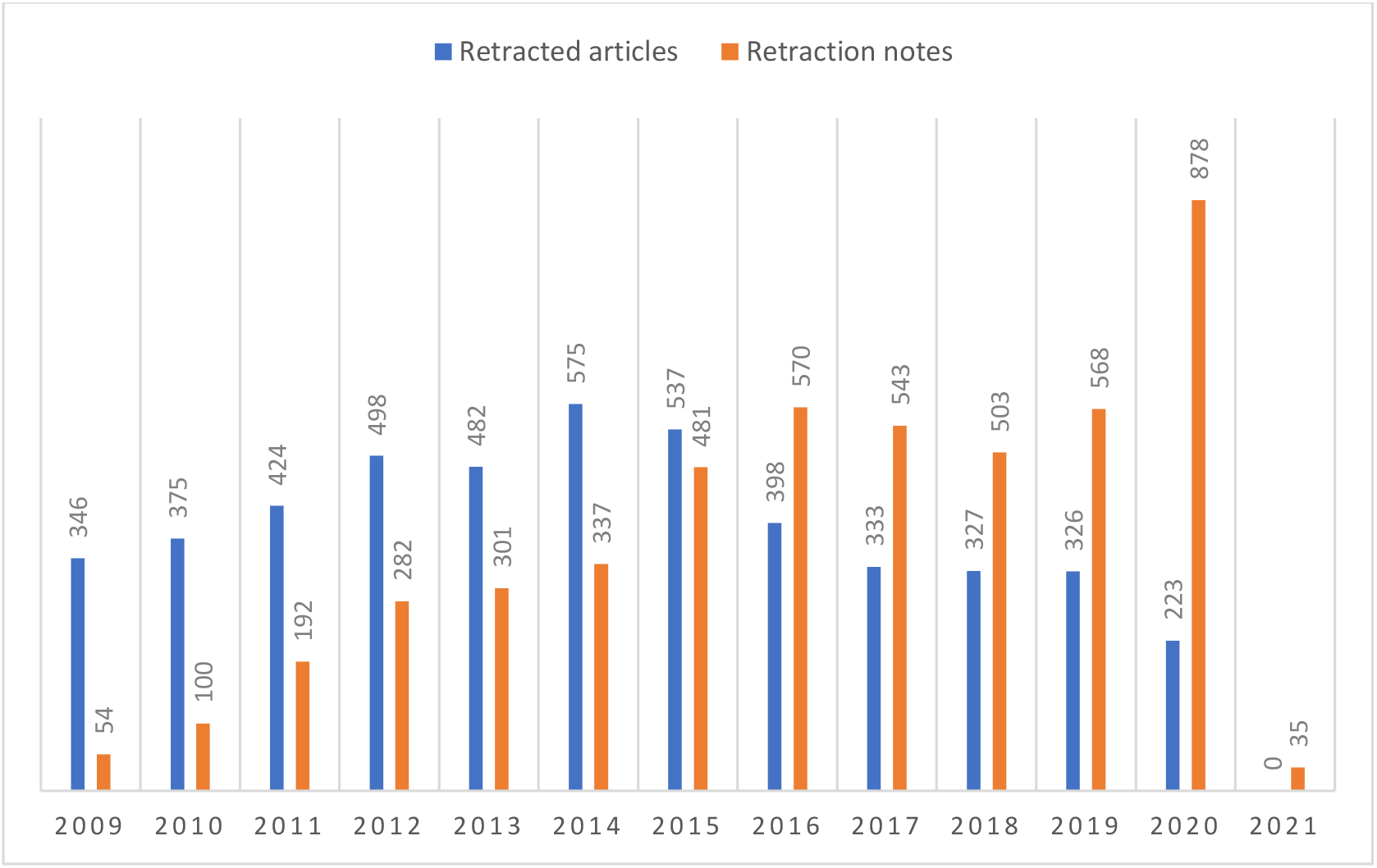
Retracted articles and retraction notes.

**Table 8.** Retracted articles and retraction notes by year (*based on PubMed publication year value and PubMed reported retraction note date; 20 articles present date errors because the date of the electronic publication was far ahead of the print publication/journal date. These errors are not reflected in the E. T. calculations.*)

|  | R.N. Year |  |  |  |  |  |  |  |  |  |  |  |  | Total |
| --- | --- | --- | --- | --- | --- | --- | --- | --- | --- | --- | --- | --- | --- | --- |
|  | 2009 | 2010 | 2011 | 2012 | 2013 | 2014 | 2015 | 2016 | 2017 | 2018 | 2019 | 2020 | 2021 |  |
| 2009 | 54 | 52 | 58 | 32 | 31 | 32 | 9 | 24 | 10 | 14 | 11 | 16 | 3 | 346 |
| 2010 | 0 | 45 | 87 | 52 | 38 | 25 | 25 | 33 | 12 | 19 | 13 | 25 | 1 | 375 |
| 2011 | 0 | 0 | 50 | 108 | 52 | 30 | 37 | 37 | 26 | 31 | 31 | 21 | 1 | 424 |
| 2012 | 0 | 0 | 0 | 90 | 110 | 54 | 45 | 59 | 42 | 27 | 35 | 34 | 2 | 498 |
| 2013 | 0 | 0 | 0 | 0 | 66 | 118 | 75 | 53 | 44 | 33 | 40 | 50 | 3 | 482 |
| 2014 | 0 | 0 | 0 | 0 | 2 | 76 | 175 | 65 | 92 | 49 | 57 | 59 | 0 | 575 |
| 2015 | 0 | 0 | 0 | 0 | 0 | 2 | 112 | 171 | 113 | 59 | 38 | 42 | 0 | 537 |
| 2016 | 0 | 0 | 0 | 0 | 0 | 0 | 3 | 125 | 111 | 54 | 48 | 54 | 3 | 398 |
| 2017 | 0 | 0 | 0 | 0 | 0 | 0 | 0 | 3 | 92 | 103 | 67 | 64 | 4 | 333 |
| 2018 | 0 | 0 | 0 | 0 | 0 | 0 | 0 | 0 | 1 | 108 | 113 | 101 | 4 | 327 |
| 2019 | 0 | 0 | 0 | 0 | 1 | 0 | 0 | 0 | 1 | 4 | 114 | 200 | 6 | 326 |
| 2020 | 0 | 0 | 0 | 0 | 0 | 0 | 0 | 0 | 0 | 0 | 3 | 212 | 8 | 223 |
| Total | 54 | 97 | 195 | 282 | 300 | 337 | 481 | 570 | 544 | 501 | 570 | 878 | 35 | 4844 |

### Retraction reasons

Out of the 4844 analyzed retractions, 4251 (87,76%) have a unique retraction reason, and 593(12,24%) have multiple retraction reasons.

**Table 9.** Retraction reasons(Column 1 displays the number of papers with a single reason as the basis for retraction. Columns 2,3,4 contain the number of papers with 2,3 and 4 concurrent reasons as the basis for retraction)

|  | Retraction Reason | All | 1 | 2 | 3 | 4 | % of 4844 papers |
| --- | --- | --- | --- | --- | --- | --- | --- |
| 1 | Mistakes/inconsistent data | 1553 | **1247** | 274 | 31 | 1 | 32,06% |
| 2 | Images | 1088 | **887** | 188 | 13 | - | 22,46% |
| 3 | Plagiarism | 663 | **519** | 115 | 27 | 2 | 13,69% |
| 4 | Overlap | 556 | **441** | 94 | 19 | 2 | 11,48% |
| 5 | Fraud | 393 | **317** | 51 | 24 | 1 | 8,11% |
| 6 | Ethics | 360 | **196** | 147 | 17 | - | 7,43% |
| 7 | Authorship | 281 | **83** | 147 | 49 | 2 | 5,78% |
| 8 | Unclear | 247 | **246** | 1 | - | - | 5,1% |
| 9 | Editor | 181 | **176** | 5 | - | - | 3,73% |
| 10 | Property & Legal | 121 | **84** | 29 | 7 | 1 | 2,5% |
| 11 | Other | 63 | **55** | 7 | 1 | - | 1,30% |
|  |  |  | **4251** |  |  |  |  |

There were 229 instances of data fabrication, 217 in the “Mistakes/Inconsistent data” category, and 12 in other categories.

We have identified 286 instances of duplicate publication, 123 were editorial/publisher mistakes, and 163 were duplicate submissions.

Within the Images category, we found 253 instances of image overlap and 94 instances of plagiarized images. When taking into account images overlap and plagiarism, the total number of overlap articles(text and images) is 809 and the total number of plagiarism articles (text and images) is 757. Image reasons count 741 cases when image overlap and plagiarism are removed.

Fraudulent peer review was found in 350 instances (main category: Fraud)

The authors were unable to provide the raw data or the raw data that could not be retrieved in 293 cases.

**Table 10.** Secondary retraction reasons

| Secondary retraction reasons | N |
| --- | --- |
| CONFLICT OF INTEREST | 31 |
| DUPLICATE (submission by authors or editorial error) | 286 |
| FABRICATED DATA | 229 |
| FALSIFIED INFORMATIONS | 39 |
| FRAUDULENT PEER REVIEW | 350 |
| NO DATA PROVIDED | 293 |
| NO IRB APPROVAL | 134 |
| NO PATIENT CONSENT | 22 |
| PUBLICATION W/O APPROVAL | 97 |
| REPRODUCIBILITY | 180 |
| RESEARCH MISCONDUCT | 47 |
| RESEARCH OUTSOURCING/TEMPLATED | 22 |
| OTHER | 14 |

### Citations

Citations were counted for all retracted papers. Overall, 140810 citations were retrieved in Google Scholar and 96000 in Dimensions (68% ratio Dimensions/Google Scholar).

**Table 11.** Citations for all 4844 retracted papers.

|  | Mean(95% CI) | Median | SEM | SD | Range | Sum |
| --- | --- | --- | --- | --- | --- | --- |
| **Google Scholar** | 29,07(26,43-31,71) | 11 | 1,346 | 93,659 | 4482 | 140810 |
| **Dimensions** | 19,82(18,08-21,55) | 7 | 0,884 | 61,547 | 2860 | 96000 |

### Exposure time

Exposure time(time difference between the most recent retraction date and the earliest publication date, expressed in months) was collected for all articles included in the study(table 11). The average exposure time was 28,89 months with a median value of 19 months.

**Table 11.** Exposure time for all 4844 retracted papers.

|  | Mean(95% CI) | Median | SEM | SD | Range |
| --- | --- | --- | --- | --- | --- |
| Exposure time(months) | 28,89(28,10-29,68) | 19 | 0,403 | 28,052 | 155 |

### Retraction reasons, exposure time, average citations/article, average number of authors and average CiteScore

**Table 12.**
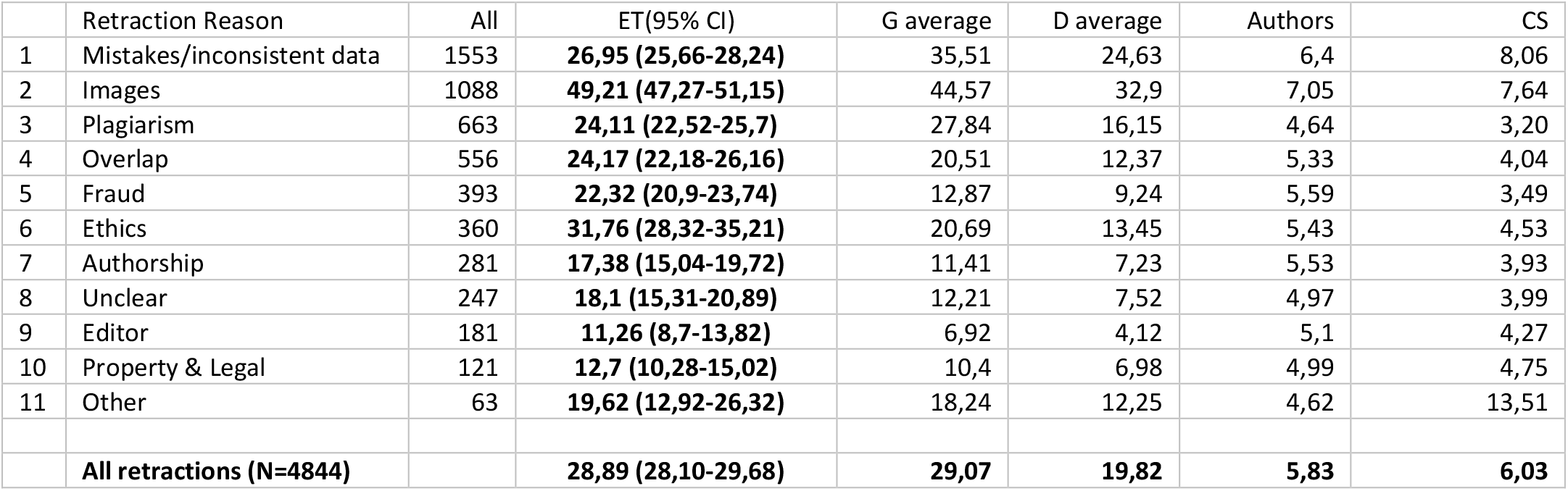
Main retraction reason categories(all instances): exposure time(E.T.), average Google Scholar citations), average Dimensions citations, the average number of authors, and the average value of CiteScore(C.S.)

**Table 13.** Main retraction reasons (papers with one retraction reason only)

|  | Retraction Reason | Single instances | ET(95% CI) | G average | D average | Authors | CS |
| --- | --- | --- | --- | --- | --- | --- | --- |
| 1 | Mistakes/inconsistent data | 1247 | **25,24 (23,89-26,6)** | 37,16 | 25,88 | 6,31 | 8,51 |
| 2 | Images | 887 | **52,01 (49,83-54,19)** | 47,29 | 34,96 | 7,16 | 7,67 |
| 3 | Plagiarism | 519 | **22,38 (20,64-24,12)** | 29,77 | 16,67 | 4,10 | 3,04 |
| 4 | Overlap | 441 | **22,74 (20,58-24,89)** | 20,02 | 11,49 | 5,21 | 3,72 |
| 5 | Fraud | 317 | **22,83 (21,29-24,38)** | 12,56 | 9,11 | 5,21 | 3,37 |
| 6 | Ethics | 196 | **32,35 (27,56-37,13)** | 18,32 | 11,6 | 5,50 | 4,33 |
| 7 | Authorship | 83 | **14,14 (10,15-18,14)** | 9,24 | 5,75 | 5,12 | 3,88 |
| 8 | Unclear | 246 | **18,14 (15,34-20,94)** | 12,26 | 7,55 | 4,96 | 4 |
| 9 | Editor | 176 | **11,45 (8,83 – 14,08)** | 7,08 | 4,20 | 5,14 | 4,35 |
| 10 | Property & Legal | 84 | **13 (10,16-15,94)** | 10,35 | 6,95 | 5,51 | 4,98 |
| 11 | Other | 55 | **19,36 (12,06-26,67)** | 18,98 | 13 | 3,73 | 14,55 |

### Number of authors

The average number of authors for the 4844 articles is 5.83(5,73-5,93) with a median of 5, mode 4, IQR 4, and range 36. The distribution of articles according to the number of authors can is displayed in Figure 3

**Figure 3.**
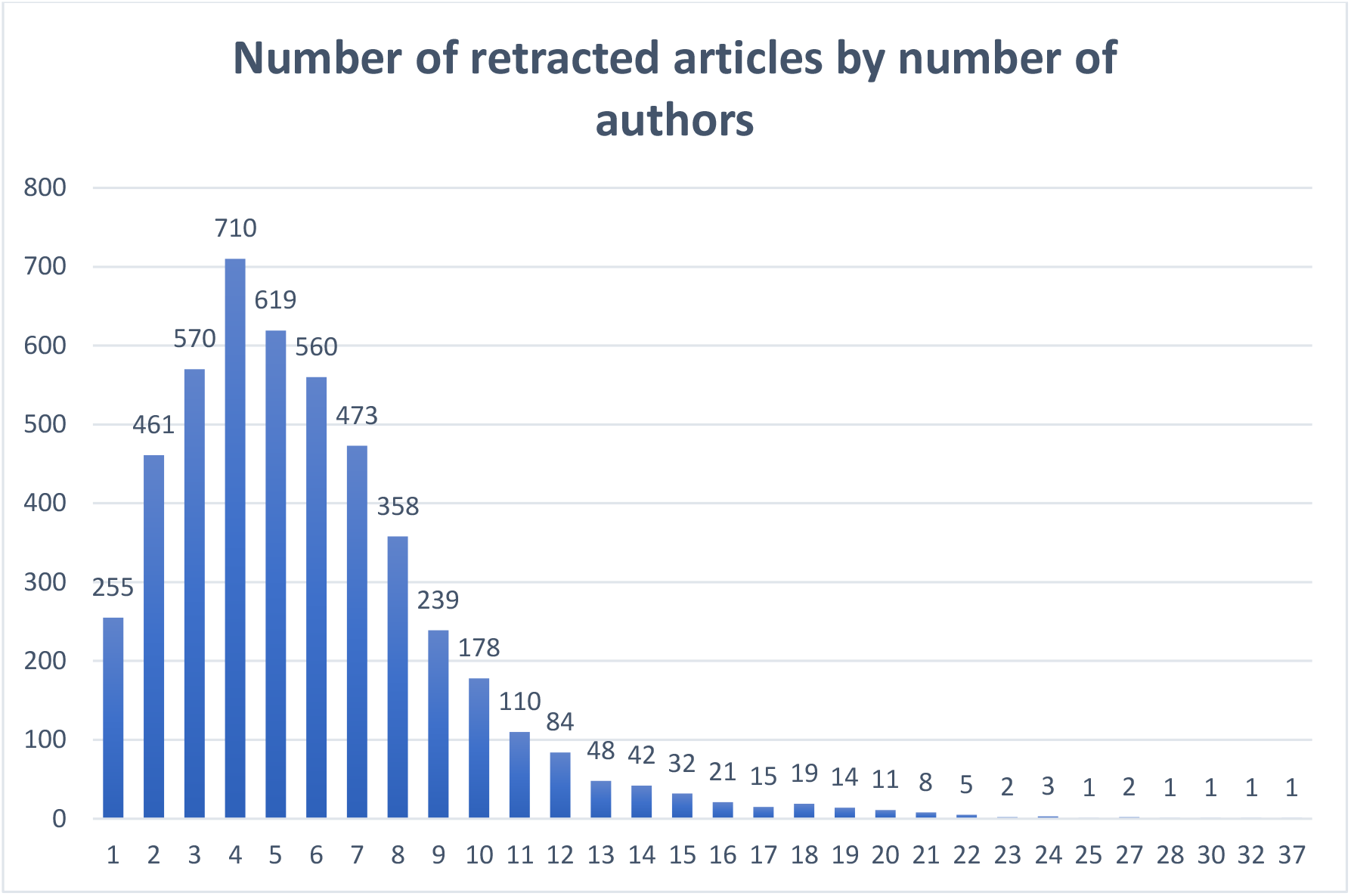
The number of retracted papers(n=4844) by the number of authors.

**Figure 4.**
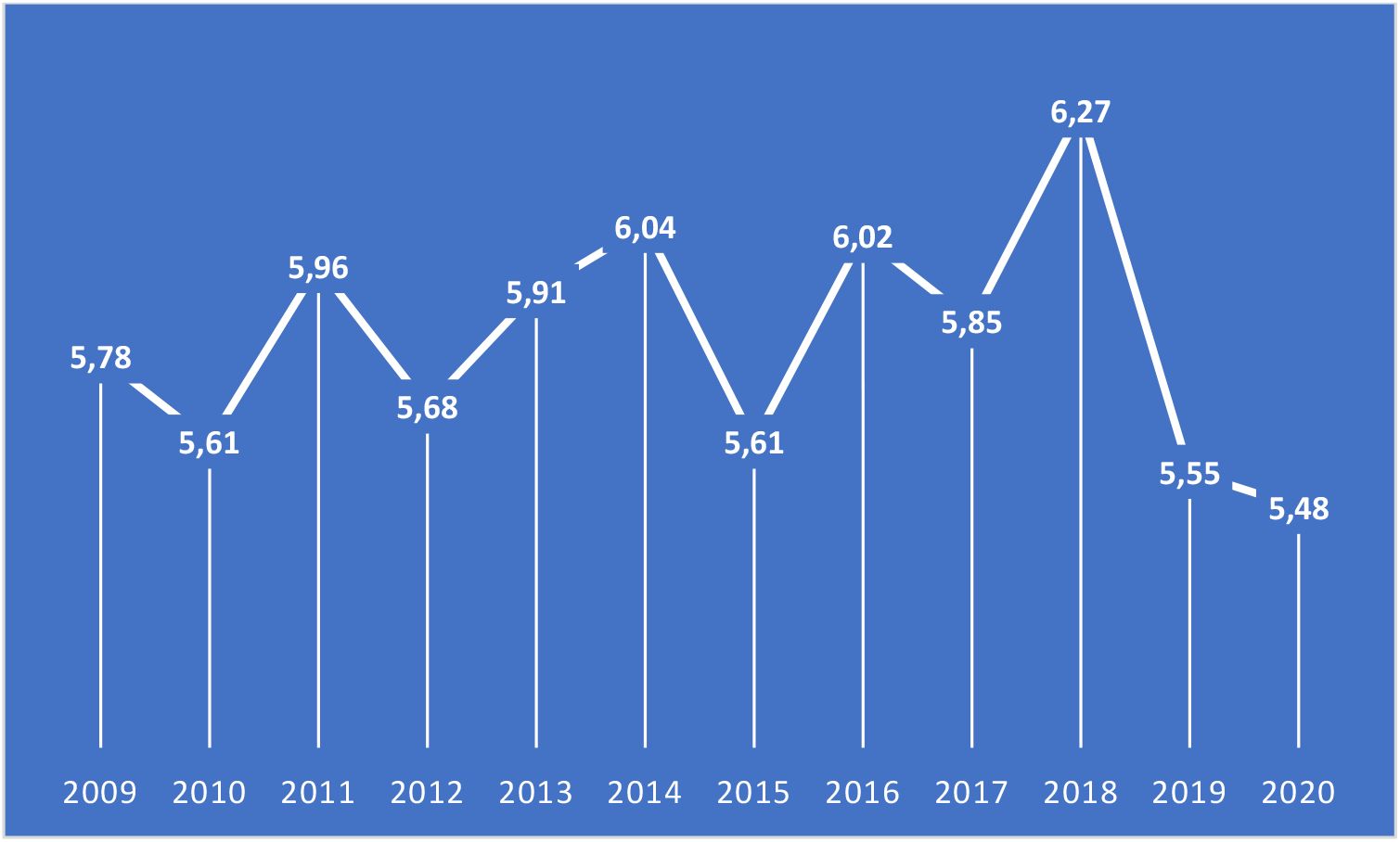
The average number of authors by year.

Of the 4844 articles analyzed, 4109 were published by authors from the same country and 735 by authors from 2 or more countries. Two hundred twenty-five (5.3%) articles had only one author, 4589 (94.7%) had two or more authors.

The average number of authors varied between 5,48(2020) and 6,27(2018).

### Retracted articles and retraction notes by country of the first author

The 4844 articles had the first authors from 94 countries. The top 30 countries have 4592 retractions (94.79% of total retractions).

The number and evolution of retractions and retraction notes for the top 30 countries are presented in table 14.

**Table 14.** Yearly distribution of retracted papers(R) & retraction notes(R.N.) by country of the first author.

|  |  | T | 2009 | 2010 | 2011 | 2012 | 2013 | 2014 | 2015 | 2016 | 2017 | 2018 | 2019 | 2020 | 2021 |
| --- | --- | --- | --- | --- | --- | --- | --- | --- | --- | --- | --- | --- | --- | --- | --- |
| **China** | R | 1588 | 41 | 40 | 43 | 67 | 147 | 256 | 267 | 159 | 142 | 136 | 160 | 130 |  |
|  | RN |  | 15 | 16 | 29 | 30 | 46 | 65 | 204 | 169 | 234 | 138 | 203 | 423 | 16 |
| **United States** | R | 913 | 101 | 106 | 147 | 118 | 85 | 79 | 73 | 53 | 46 | 39 | 38 | 28 |  |
|  | RN |  | 14 | 18 | 45 | 52 | 49 | 59 | 107 | 122 | 103 | 109 | 96 | 135 | 4 |
| **India** | R | 351 | 23 | 30 | 38 | 51 | 42 | 53 | 37 | 25 | 22 | 12 | 13 | 5 |  |
|  | RN |  | 2 | 3 | 21 | 31 | 36 | 32 | 32 | 35 | 22 | 35 | 48 | 51 | 3 |
| **Japan** | R | 212 | 35 | 34 | 31 | 26 | 12 | 14 | 14 | 15 | 9 | 11 | 7 | 5 |  |
|  | RN |  | 4 | 6 | 19 | 17 | 20 | 17 | 15 | 17 | 12 | 31 | 20 | 34 | 1 |
| **Italy** | R | 182 | 12 | 20 | 15 | 32 | 36 | 21 | 11 | 10 | 4 | 9 | 8 | 4 |  |
|  | RN |  | 0 | 4 | 8 | 15 | 8 | 24 | 12 | 13 | 18 | 20 | 35 | 23 | 2 |
| **South Korea** | R | 166 | 7 | 13 | 31 | 22 | 18 | 9 | 10 | 14 | 11 | 15 | 9 | 7 |  |
|  | RN |  | 2 | 2 | 2 | 46 | 15 | 8 | 9 | 15 | 10 | 13 | 13 | 31 | 0 |
| **Iran** | R | 162 | 6 | 10 | 6 | 16 | 16 | 25 | 31 | 17 | 11 | 11 | 9 | 4 |  |
|  | RN |  | 1 | 4 | 4 | 2 | 10 | 9 | 18 | 57 | 10 | 9 | 22 | 15 | 1 |
| **Germany** | R | 115 | 29 | 17 | 14 | 11 | 6 | 5 | 8 | 7 | 3 | 5 | 4 | 6 |  |
|  | RN |  | 1 | 7 | 21 | 15 | 8 | 8 | 11 | 8 | 12 | 0 | 9 | 14 | 1 |
| **United Kingdom** | R | 111 | 13 | 13 | 7 | 14 | 9 | 13 | 9 | 6 | 8 | 13 | 3 | 3 |  |
|  | RN |  | 2 | 4 | 7 | 10 | 9 | 8 | 9 | 11 | 11 | 14 | 9 | 17 | 0 |
| **Spain** | R | 79 | 6 | 3 | 5 | 15 | 6 | 5 | 13 | 8 | 3 | 6 | 2 | 7 |  |
|  | RN |  | 1 | 1 | 2 | 5 | 4 | 3 | 3 | 13 | 8 | 19 | 5 | 14 | 1 |
| **Canada** | R | 78 | 7 | 9 | 8 | 8 | 8 | 4 | 6 | 7 | 8 | 4 | 6 | 3 |  |
|  | RN |  | 0 | 3 | 2 | 5 | 7 | 9 | 5 | 13 | 6 | 9 | 6 | 11 | 2 |
| **France** | R | 59 | 3 | 7 | 3 | 7 | 6 | 4 | 2 | 3 | 9 | 7 | 5 | 2 |  |
|  | RN |  | 0 | 1 | 4 | 2 | 6 | 6 | 2 | 5 | 4 | 11 | 7 | 10 | 0 |
| **Egypt** | R | 55 | 3 | 7 | 5 | 7 | 6 | 7 | 3 | 6 | 2 | 8 | 3 | 0 |  |
|  | RN |  | 0 | 1 | 6 | 2 | 9 | 6 | 2 | 6 | 4 | 7 | 8 | 6 | 0 |
| **Australia** | R | 55 | 7 | 10 | 1 | 6 | 4 | 5 | 1 | 6 | 3 | 2 | 8 | 2 |  |
|  | RN |  | 3 | 6 | 4 | 2 | 3 | 3 | 3 | 8 | 6 | 6 | 5 | 6 | 0 |
| **Sweden** | R | 54 | 4 | 3 | 5 | 4 | 10 | 8 | 3 | 4 | 3 | 3 | 4 | 3 |  |
|  | RN |  | 1 | 2 | 1 | 2 | 3 | 6 | 5 | 3 | 5 | 10 | 8 | 8 | 0 |
| **Turkey** | R | 53 | 2 | 4 | 4 | 4 | 5 | 7 | 9 | 6 | 3 | 5 | 3 | 1 |  |
|  | RN |  | 1 | 2 | 1 | 5 | 4 | 7 | 8 | 6 | 7 | 3 | 4 | 5 | 0 |
| **Brazil** | R | 51 | 3 | 5 | 5 | 14 | 6 | 3 | 5 | 2 | 2 | 2 | 2 | 2 |  |
|  | RN |  | 0 | 0 | 0 | 2 | 8 | 4 | 1 | 17 | 6 | 4 | 4 | 5 | 0 |
| **Taiwan** | R | 49 | 2 | 2 | 2 | 6 | 9 | 7 | 5 | 7 | 1 | 4 | 3 | 1 |  |
|  | RN |  | 0 | 0 | 1 | 1 | 4 | 2 | 4 | 7 | 11 | 5 | 5 | 8 | 1 |
| **Netherlands** | R | 40 | 4 | 5 | 12 | 2 | 2 | 2 | 1 | 4 | 3 | 1 | 3 | 1 |  |
|  | RN |  | 0 | 0 | 2 | 9 | 5 | 4 | 2 | 5 | 4 | 2 | 3 | 3 | 1 |
| Saudi Arabia | R | 36 | 1 | 2 | 5 | 2 | 5 | 2 | 3 | 0 | 3 | 9 | 2 | 1 |  |
|  | RN |  | 1 | 1 | 1 | 1 | 1 | 4 | 1 | 3 | 2 | 5 | 7 | 6 | 2 |
| Singapore | R | 31 | 4 | 4 | 3 | 6 | 2 | 3 | 2 | 4 | 2 | 0 | 1 | 0 |  |
|  | RN |  | 0 | 1 | 2 | 0 | 3 | 4 | 3 | 5 | 1 | 5 | 3 | 4 | 0 |
| Switzerland | R | 28 | 4 | 1 | 3 | 7 | 3 | 2 | 4 | 2 | 0 | 0 | 1 | 1 |  |
|  | RN |  | 2 | 0 | 0 | 1 | 7 | 5 | 1 | 1 | 6 | 3 | 1 | 1 | 0 |
| Poland | R | 18 | 0 | 0 | 0 | 1 | 2 | 3 | 2 | 6 | 2 | 2 | 0 | 0 |  |
|  | RN |  | 0 | 0 | 0 | 0 | 0 | 1 | 3 | 1 | 4 | 3 | 5 | 1 | 0 |
| Pakistan | R | 18 | 2 | 1 | 0 | 2 | 1 | 0 | 0 | 2 | 2 | 4 | 2 | 2 |  |
|  | RN |  | 1 | 1 | 1 | 2 | 2 | 0 | 0 | 1 | 2 | 1 | 4 | 3 | 0 |
| Israel | R | 16 | 0 | 2 | 3 | 3 | 1 | 3 | 0 | 1 | 0 | 1 | 2 | 0 |  |
|  | RN |  | 0 | 0 | 2 | 2 | 1 | 0 | 1 | 0 | 3 | 3 | 3 | 1 | 0 |
| Malaysia | R | 15 | 0 | 1 | 0 | 3 | 1 | 3 | 1 | 5 | 1 | 0 | 0 | 0 |  |
|  | RN |  | 0 | 0 | 1 | 1 | 2 | 0 | 0 | 3 | 2 | 5 | 0 | 1 | 0 |
| Greece | R | 15 | 3 | 0 | 2 | 2 | 1 | 3 | 1 | 2 | 0 | 1 | 0 | 0 |  |
|  | RN |  | 1 | 0 | 1 | 0 | 2 | 2 | 2 | 2 | 3 | 3 | 3 | 1 | 0 |
| Russian Federation | R | 14 | 0 | 0 | 1 | 2 | 1 | 4 | 2 | 1 | 3 | 0 | 0 | 0 |  |
|  | RN |  | 0 | 0 | 0 | 0 | 2 | 0 | 0 | 0 | 6 | 1 | 4 | 1 | 0 |
| Norway | R | 14 | 0 | 1 | 1 | 1 | 3 | 2 | 0 | 1 | 2 | 2 | 0 | 1 |  |
|  | RN |  | 0 | 0 | 0 | 1 | 1 | 0 | 1 | 0 | 0 | 3 | 3 | 5 | 0 |
| Ireland | R | 14 | 1 | 2 | 2 | 3 | 2 | 2 | 0 | 1 | 1 | 0 | 0 | 0 |  |
|  | RN |  | 0 | 1 | 0 | 0 | 1 | 3 | 0 | 1 | 3 | 4 | 1 | 0 | 0 |

### Retraction reasons for top 10 countries, first author country

**Table 15.** Retraction reasons for top 10 countries (first author affiliation considered).

| Country | N | MISTAKES | IMAGES | PLAGIARISM | OVERLAP | FRAUD | ETHICS | AUTHORSHIP | UNCLEAR | EDITOR | PROPERTY | OTHER |
| --- | --- | --- | --- | --- | --- | --- | --- | --- | --- | --- | --- | --- |
| China | 1588 | 28,6% | 23,2% | 14,7% | 9,3% | 18,3% | 5,7% | 6,6% | 5% | 2% | 1,4% | 0,5% |
| United States | 913 | 45,3% | 31,2% | 3,8% | 7,1% | 0,5% | 5,5% | 1,8% | 3,8% | 4,2% | 2,8% | 1,5% |
| India | 351 | 17,9% | 24,8% | 35,3% | 17,1% | 1,1% | 5,1% | 4,3% | 5,7% | 2,6% | 1,1% | 0,3% |
| Japan | 212 | 39,2% | 19,8% | 4,7% | 16,5% | 0,5% | 15,6% | 9% | 5,7% | 3,8% | 2,8% | 0,5% |
| Italy | 182 | 12,1% | 27,5% | 32,4% | 12,1% | 0,5% | 6% | 3,3% | 5,5% | 4,4% | 3,8% | 2,7% |
| South Korea | 166 | 33,1% | 10,8% | 6,6% | 15,1% | 16,9% | 12% | 4,8% | 4,8% | 2,4% | 4,2% | 0,6% |
| Iran | 162 | 14,2% | 4,9% | 29,6% | 22,2% | 32,7% | 4,9% | 23,5% | 5,6% | 5,6% | 1,2% | 0,6% |
| Germany | 115 | 36,5% | 7,8% | 1,7% | 15,7% | 0 | 20,9% | 7% | 4,3% | 6,1% | 5,2% | 2,6% |
| United Kingdom | 111 | 39,6% | 20,7% | 5,4% | 6,3% | 0 | 9% | 9% | 4,5% | 8,1% | 5,4% | 3,6% |
| Spain | 79 | 26,6% | 36,7% | 8,9% | 12,7% | 1,3% | 3,8% | 5,1% | 5,1% | 3,8% | 6,3% | 1,3% |

### Retraction reasons for top 10 countries, all authors countries

**Table 16.** Retraction reasons for top 10 countries (all author countries).

| Country | N | MISTAKES | IMAGES | PLAGIARISM | OVERLAP | FRAUD | ETHICS | AUTHORSHIP | UNCLEAR | EDITOR | PROPERTY | OTHER |
| --- | --- | --- | --- | --- | --- | --- | --- | --- | --- | --- | --- | --- |
| China | 1626 | 29% | 23,5% | 14,4% | 9,4% | 18% | 5,8% | 6,6% | 4,9% | 1,9% | 1,5% | 0,5% |
| United States | 1175 | 43,2% | 31,4% | 4,9% | 7,6% | 1,4% | 6,1% | 2,7% | 3,7% | 4,2% | 2,6% | 1,7% |
| India | 384 | 18,2% | 27,1% | 33,6% | 16,1% | 1,3% | 4,9% | 4,7% | 5,7% | 2,3% | 1,3% | 0,3% |
| Japan | 241 | 40,2% | 20,7% | 5% | 14,9% | 1,2% | 14,1% | 7,9% | 5,8% | 3,3% | 2,5% | 0,4% |
| Italy | 227 | 21,6% | 29,1% | 26% | 11% | 0,9% | 5,7% | 2,6% | 4,4% | 4% | 3,5% | 2,2% |
| United Kingdom | 194 | 44,1% | 18,5% | 6,7% | 7,2% | 0,5% | 10,8% | 8,2% | 2,6% | 5,1% | 4,1% | 4,1% |
| South Korea | 182 | 32,4% | 14,8% | 7,7% | 13,7% | 15,4% | 11% | 4,4% | 4,4% | 2,2% | 3,8% | 0,5% |
| Germany | 177 | 37,3% | 16,4% | 2,8% | 12,4% | 1,7% | 16,4% | 9% | 2,8% | 5,1% | 5,1% | 4% |
| Iran | 175 | 14,9% | 6,9% | 27,4% | 21,7% | 30,9% | 6,3% | 22,9% | 6,3% | 5,1% | 1,1% | 0,6% |
| Canada | 109 | 32,1% | 20,2% | 4,6% | 14,7% | 2,8% | 8,3% | 5,5% | 2,8% | 11% | 3,7% | 5,5% |

The distribution of retraction reasons is not uniform and seems to reflect country-specific problems.

### Impact of retracted articles

In order to estimate the impact of retracted research, we calculated for each country the number of articles and percentage from total retractions(%A), the number of Google Scholar citations and percentage from all citations(%G), the number of Dimensions citations and percentage from all Dimensions citations(%D), the ET and CiteScore average.

Seven categories were defined: four marked green (positive or stationary evolution) and three red (negative evolution).

The high impact was considered any difference between %G or %D and %A greater than 25% (with steps at >25, >50, and >75%).

Low impact was considered any difference between%G or %D and %A less than 25%.

Values between −25% and + 25% were considered stationary/neutral.

The impact assessment was made for:

- first author country of origin(with and without editorial errors)
- all authors countries of origin(with and without editorial errors)

**Table 17.**
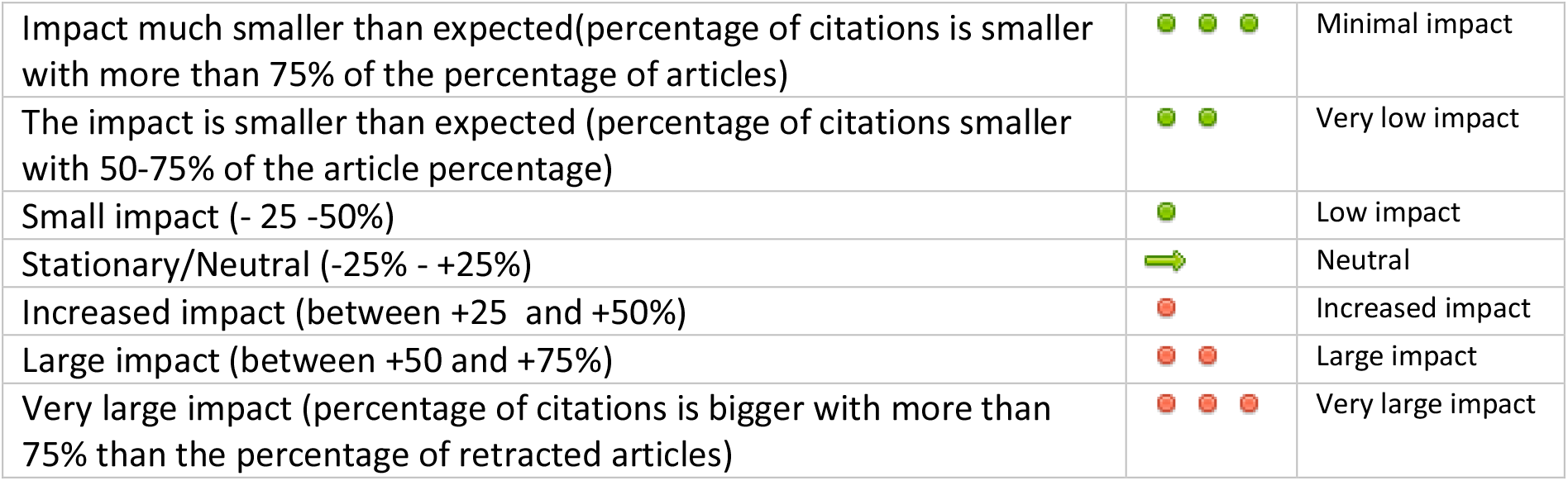
Impact grading.

|  |  |  |
| --- | --- | --- |
| Impact much smaller than expected(percentage of citations is smaller with more than 75% of the percentage of articles) |  | Minimal impact |
| The impact is smaller than expected (percentage of citations smaller with 50-75% of the article percentage) |  | Very low impact |
| Small impact (- 25 -50%) |  | Low impact |
| Stationary/Neutral (-25% - +25%) |  | Neutral |
| Increased impact (between +25 and +50%) |  | Increased impact |
| Large impact (between +50 and +75%) |  | Large impact |
| Very large impact (percentage of citations is bigger with more than 75% than the percentage of retracted articles) |  | Very large impact |

## Discussions

### Trend

The number of articles withdrawn from the biomedical literature is increasing, a fact that was constantly signaled in the articles that study this subject[26,28,37,38,43–45].

We have not identified elements to signal a slowdown. On the contrary, 2020 seems to be a record year for retraction notes, 878 (18.6% of the total) being already registered in PubMed on January 31st, 2021. Almost half of the 2020 retractions(423/878) are issued for articles with the first author affiliated to a Chinese institution and 135 for authors affiliated to US institutions. Considering the period 2009-2020, five of the top 10 countries recorded the highest number of withdrawals in 2020: China, United States, India, Japan, and the United Kingdom.

The process of correcting the biomedical literature seems to be now continuous and consistent, going back ten years or more, 11% of the retraction notes appearing in 2020 and 15,8% in 2019, being for papers published in 2009-2012(see table 8).

### Countries

More than 50% of the total number of retracted articles come from China (1588 / 32.78%) and the United States (913 / 18,84%), followed by India (351 / 7,24%), Japan (213 / 4,37%) and Italy (182 / 3,75%) (complete information for the first 30 countries is in table 18).

**Table 18.** Retracted research impact for top 30 countries(first author, all retraction reasons). 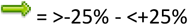; 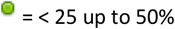; 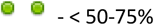; 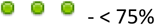; 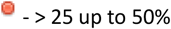; 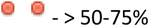; 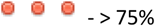; CS – CiteScore average(3933 records with available information) ; ET – Exposure Time average

|  |  | N | %A | G | %G |  | D | %D |  | CS | ET | Authors |
| --- | --- | --- | --- | --- | --- | --- | --- | --- | --- | --- | --- | --- |
| 1 | China | 1588 | 32,78% | 23011 | 16,34% | ● ● | 17245 | 17,96% | ● | 4,4 | 22,93 | 6,2 |
| 2 | United States | 913 | 18,84% | 48824 | 34,67% | ● ● ● | 33947 | 35,36% | ● ● ● | 9,94 | 39,16 | 5,8 |
| 3 | India | 351 | 7,24% | 13231 | 9,39% | ● | 7708 | 8,03% | ➡ | 4,38 | 31,74 | 4,5 |
| 4 | Japan | 212 | 4,37% | 5477 | 3,89% | ➡ | 3645 | 3,79% | ➡ | 5,67 | 39,24 | 7,1 |
| 5 | Italy | 182 | 3,75% | 4821 | 3,43% | ➡ | 3283 | 3,42% | ➡ | 4,93 | 38,14 | 6,4 |
| 6 | South Korea | 166 | 3,42 | 2606 | 1,85% | ● | 1718 | 1,79% | ● | 4,44 | 21,61 | 5,3 |
| 7 | Iran | 162 | 3,34 | 3346 | 2,37% | ● | 2072 | 2,16% | ● | 3,34 | 23,04 | 5 |
| 8 | Germany | 115 | 2,2% | 3100 | 2,2% | ➡ | 2044 | 2,13% | ➡ | 5,78 | 26,96 | 5,6 |
| 9 | United Kingdom | 111 | 2,29% | 5478 | 3,89% | ● ● | 3908 | 4,07% | ● ● ● | 13,59 | 27,1 | 4,7 |
| 10 | Spain | 79 | 1,63% | 7515 | 5,33% | ● ● ● | 5040 | 5,25% | ● ● ● | 8,89 | 33,54 | 6,6 |
| 11 | Canada | 78 | 1,61% | 1584 | 1,12% | ● | 1059 | 1,1% | ● | 7,21 | 28,18 | 5,6 |
| 12 | France | 59 | 1,22% | 1785 | 1,27% | ➡ | 1313 | 1,36% | ➡ | 10,44 | 24,29 | 6,7 |
| 13 | Egypt | 55 | 1,13% | 2410 | 1,71% | ● ● | 1476 | 1,54% | ● | 4,64 | 25,8 | 4,2 |
| 14 | Australia* | 55 | 1,13% | 1018 | 0,72% | ● | 1050 | 1,09% | ➡ | 6,28 | 19,25 | 4,4 |
| 15 | Sweden | 54 | 1,11% | 1795 | 1,27% | ➡ | 1346 | 1,4% | ● | 10,43 | 29,74 | 6,6 |
| 16 | Turkey | 53 | 1,09% | 419 | 0,29% | ● ● ● | 202 | 0,21% | ● ● ● | 2,85 | 14,94 | 4,9 |
| 17 | Brazil | 51 | 1,05% | 1980 | 1,4% | ● | 1268 | 1,32% | ● | 5,3 | 36,18 | 6,8 |
| 18 | Taiwan | 49 | 1,01% | 1037 | 0,73% | ● | 728 | 0,76% | ➡ | 8,13 | 35,76 | 6,3 |
| 19 | Netherlands | 40 | 0,83% | 2043 | 1,45% | ● ● | 967 | 1% | ➡ | 8,57 | 26,23 | 5,8 |
| 20 | Saudi Arabia | 36 | 0,74% | 593 | 0,42% | ● | 424 | 0,44 | ● | 3,61 | 29,14 | 4,1 |
| 21 | Singapore | 31 | 0,64% | 1410 | 1% | ● ● | 1027 | 1,07% | ● ● | 8,28 | 40,58 | 5,9 |
| 22 | Switzerland | 28 | 0,58% | 973 | 0,69% | ➡ | 625 | 0,65% | ➡ | 13,66 | 25,14 | 6,5 |
| 23 | Pakistan | 18 | 0,37% | 73 | 0,05% | ● ● ● | 49 | 0,05% | ● ● ● | 3,7 | 7,06 | 5,7 |
| 24 | Poland | 18 | 0,37% | 261 | 0,18% | ● ● | 190 | 0,2% | ● | 4,85 | 23,94 | 6,7 |
| 25 | Israel | 16 | 0,33% | 402 | 0,28% | ➡ | 285 | 0,29% | ➡ | 8,95 | 31,5 | 8,8 |
| 26 | Greece | 15 | 0,31% | 236 | 0,17% | ● | 125 | 0,13% | ● ● | 7,35 | 24,8 | 4,5 |
| 27 | Malaysia | 15 | 0,31% | 195 | 0,14% | ● ● | 125 | 0,13% | ● ● | 3,08 | 21,8 | 5,7 |
| 28 | Ireland | 14 | 0,29% | 545 | 0,39% | ● | 344 | 0,36% | ➡ | 7,05 | 46 | 4,8 |
| 29 | Norway | 14 | 0,29% | 377 | 0,27% | ➡ | 251 | 0,26% | ➡ | 4,7 | 40,57 | 7,1 |
| 30 | Russian Federation | 14 | 0,29% | 211 | 0,15% | ● | 111 | 0,11% | ● ● | 4,13 | 35,21 | 5,5 |
|  | Totals/averages | 4844 |  | 140810 |  |  | 96000 |  |  | 6,04 | 28,89 | 5,83 |

**Table 19.** Retracted research impact for top 30 countries(first author, without editorial retraction reason.). 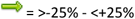; 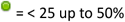; 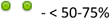; 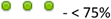; 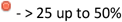; 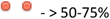; 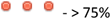; CS – CiteScore average(3933 records with available information) ; ET – Exposure Time average

|  |  | N | %A | G | %G | Evolution | D | %D | Evolution | CS | ET | Authors |
| --- | --- | --- | --- | --- | --- | --- | --- | --- | --- | --- | --- | --- |
| 1 | China | 1557 | 33,35% | 22749 | 16,3% | ● ● | 17080 | 17,93% | ● | 4,41 | 23,19 | 6,2 |
| 2 | United States | 879 | 18,83% | 48625 | 34,84% | ● ● ● | 33807 | 35,49% | ● ● ● | 10,16 | 40,33 | 5,9 |
| 3 | India | 342 | 7,32% | 13179 | 9,44% | ● | 7684 | 8,06% | ➡ | 4,4 | 31,98 | 4,5 |
| 4 | Japan | 204 | 4,37% | 5425 | 3,89% | ➡ | 3582 | 3,76% | ➡ | 5,65 | 40,67 | 7 |
| 5 | Italy | 174 | 3,73% | 4774 | 3,42% | ➡ | 3255 | 3,42% | ➡ | 4,75 | 39,28 | 6,4 |
| 6 | South Korea | 162 | 3,47% | 2586 | 1,85% | ● | 1712 | 1,8% | ● | 4,48 | 21,38 | 5,2 |
| 7 | Iran | 153 | 3,28% | 3289 | 2,35% | ● | 2028 | 2,13% | ● | 3,45 | 23,67 | 5 |
| 8 | Germany | 108 | 2,31% | 3081 | 2,21% | ➡ | 2010 | 2,11% | ➡ | 5,94 | 28,49 | 5,7 |
| 9 | United Kingdom | 102 | 2,18% | 5293 | 3,79% | ● ● | 3841 | 4,03% | ● ● ● | 13,85 | 27,16 | 4,7 |
| 10 | Spain | 77 | 1,65% | 7515 | 5,38% | ● ● ● | 5036 | 5,29% | ● ● ● | 9,07 | 34,38 | 6,7 |
| 11 | Canada | 67 | 1,43% | 1518 | 1,09% | ➡ | 1038 | 1,09% | ➡ | 7,66 | 32,07 | 5,8 |
| 12 | France | 56 | 1,2% | 1777 | 1,27% | ➡ | 1306 | 1,37% | ➡ | 10,64 | 25,09 | 6,9 |
| 13 | Egypt | 54 | 1,16% | 2409 | 1,73% | ● | 1476 | 1,55% | ● | 4,64 | 26,22 | 4,2 |
| 14 | Sweden | 51 | 1,09% | 1786 | 1,28% | ➡ | 1335 | 1,4% | ● | 10,95 | 30,18 | 6,9 |
| 15 | Australia* | 49 | 1,05% | 941 | 0,67% | ● | 1021 | 1,07% | ➡ | 6,72 | 21,16 | 4,5 |
| 16 | Turkey | 49 | 1,05% | 413 | 0,29% | ● ● | 197 | 0,2% | ● ● ● | 2,89 | 15,51 | 4,9 |
| 17 | Taiwan | 48 | 1,03% | 1036 | 0,74% | ● | 728 | 0,76% | ● | 8,27 | 36,46 | 6,3 |
| 18 | Brazil | 47 | 1% | 1945 | 1,39% | ● | 1250 | 1,31% | ● | 5,56 | 37,55 | 6,9 |
| 19 | Saudi Arabia | 36 | 0,77% | 593 | 0,42% | ● | 424 | 0,44% | ● | 3,61 | 29,14 | 4,1 |
| 20 | Netherlands | 34 | 0,73% | 2010 | 1,44% | ● ● ● | 955 | 1% | ● | 9,27 | 30 | 5,8 |
| 21 | Singapore | 31 | 0,66% | 1410 | 1,01% | ● ● | 1027 | 1,08% | ● ● | 8,28 | 40,6 | 5,9 |
| 22 | Switzerland | 28 | 0,6% | 973 | 0,7% | ➡ | 625 | 0,66% | ➡ | 13,66 | 25,14 | 6,5 |
| 23 | Poland | 18 | 0,38% | 261 | 0,19% | ● ● | 190 | 0,2% | ● | 4,85 | 23,94 | 6,7 |
| 24 | Pakistan | 17 | 0,36% | 73 | 0,05% | ● ● ● | 42 | 0,04% | ● ● ● | 3,7 | 7,35 | 5,8 |
| 25 | Israel | 16 | 0,34% | 402 | 0,29% | ➡ | 285 | 0,3% | ➡ | 8,96 | 31,5 | 8,7 |
| 26 | Greece | 15 | 0,32% | 236 | 0,17% | ● | 125 | 0,13% | ● ● | 7,36 | 24,8 | 4,5 |
| 27 | Malaysia | 15 | 0,32% | 195 | 0,14% | ● ● | 125 | 0,13% | ● ● | 3,08 | 21,8 | 5,7 |
| 28 | Ireland | 14 | 0,3% | 545 | 0,39% | ● | 344 | 0,36% | ➡ | 7,05 | 46 | 4,8 |
| 29 | Norway | 14 | 0,3% | 377 | 0,27% | ➡ | 251 | 0,26% | ➡ | 4,7 | 40,57 | 7,1 |
| 30 | Russian Federation | 14 | 0,3% | 211 | 0,15% | ● ● | 111 | 0,12% | ● ● | 4,14 | 35,21 | 5,5 |
|  | Totals/averages | 4668 |  | 139564 |  |  | 95260 |  |  | 6,1 | 29,54 | 5,85 |

All author countries and impact of retracted papers(tables 20,21).

**Table 20.** Retracted research impact for top 30 countries(all authors, all retraction reasons) 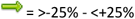; 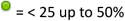; 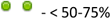; 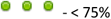; 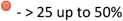; 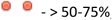; 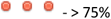; CS - CiteScore average(3933 records with available information) ; ET - Exposure Time average

|  | Country | N | %A | G | %G | Evolution | D | %D | Evolution | CS | ET | Authors |
| --- | --- | --- | --- | --- | --- | --- | --- | --- | --- | --- | --- | --- |
| 1 | China | 1626 | 33,57% | 24658 | 17,51% | ● | 18332 | 19,09% | ● | 4,56 | 23,11 | 6,3 |
| 2 | United States | 1175 | 24,26% | 58726 | 41,7% | ● ● | 40643 | 42,33% | ● ● | 9,86 | 37,1 | 6,4 |
| 3 | India | 384 | 7,93% | 15163 | 10,77% | ● | 9019 | 9,39% | ➡ | 4,41 | 32,18 | 4,7 |
| 4 | Japan | 241 | 4,97% | 8117 | 5,76% | ➡ | 5496 | 5,72% | ➡ | 6,72 | 39,87 | 7,3 |
| 5 | Italy | 227 | 4,69% | 8547 | 6,07% | ● | 5770 | 6,01% | ● | 6,37 | 39,06 | 7,7 |
| 6 | United Kingdom | 195 | 4,03% | 9781 | 6,95% | ● ● | 7355 | 7,66% | ● ● ● | 12,29 | 28,2 | 6,7 |
| 7 | South Korea | 182 | 3,76% | 3218 | 2,28% | ● | 2177 | 2,27% | ● | 4,85 | 22,96 | 5,5 |
| 8 | Germany | 177 | 3,65% | 6449 | 4,58% | ● | 4186 | 4,36% | ➡ | 8,18 | 29,55 | 7,7 |
| 9 | Iran | 175 | 3,61% | 3440 | 2,44% | ● | 2122 | 2,21% | ● | 3,4 | 22,36 | 5,1 |
| 10 | Canada | 109 | 2,25% | 2956 | 2,1% | ➡ | 1747 | 1,82% | ➡ | 7,97 | 29,18 | 6,6 |
| 11 | Spain | 100 | 2,06% | 8631 | 6,13% | ● ● ● | 5905 | 6,15% | ● ● ● | 9,32 | 34,63 | 8,1 |
| 12 | France | 95 | 1,96% | 3881 | 2,76 | ● | 2496 | 2,6% | ● | 11,2 | 29,8 | 8,3 |
| 13 | Australia | 77 | 1,59% | 1367 | 0,97% | ● | 1265 | 1,32% | ➡ | 7,03 | 18,77 | 5,5 |
| 14 | Sweden | 74 | 1,53% | 3846 | 2,73% | ● ● ● | 2634 | 2,74% | ● ● ● | 11,18 | 31,5 | 7,8 |
| 15 | Netherlands | 66 | 1,36% | 3029 | 2,15% | ● ● | 1598 | 1,66% | ➡ | 8,49 | 24,33 | 7,1 |
| 16 | Egypt | 63 | 1,3% | 2492 | 1,77% | ● | 1537 | 1,6% | ➡ | 4,45 | 25,33 | 4,3 |
| 17 | Taiwan | 59 | 1,22% | 1225 | 0,87 | ● | 864 | 0,9% | ➡ | 8,01 | 34,54 | 6,6 |
| 18 | Brazil | 57 | 1,18% | 2175 | 1,54% | ● | 1402 | 1,46% | ➡ | 5,97 | 36,96 | 6,9 |
| 19 | Turkey | 57 | 1,18% | 422 | 0,3% | ● ● | 205 | 0,21% | ● ● ● | 3,12 | 14,12 | 5,2 |
| 20 | Switzerland | 49 | 1,01% | 2145 | 1,52% | ● ● | 1397 | 1,46% | ● | 15,19 | 25,43 | 8,8 |
| 21 | Saudi Arabia | 46 | 0,95% | 1424 | 1,01% | ➡ | 793 | 0,83% | ➡ | 3,74 | 31,91 | 4,6 |
| 22 | Singapore | 39 | 0,8% | 1904 | 1,35% | ● ● | 1380 | 1,44% | ● ● ● | 8,2 | 36,62 | 6 |
| 23 | Pakistan | 28 | 0,61% | 102 | 0,07% | ● ● ● | 71 | 0,07% | ● ● ● | 3,88 | 6,39 | 5,5 |
| 24 | Norway | 26 | 0,54% | 757 | 0,54% | ➡ | 516 | 0,54% | ➡ | 6,31 | 35,12 | 8,9 |
| 25 | Greece | 25 | 0,52% | 869 | 0,62% | ➡ | 592 | 0,62% | ➡ | 8,49 | 24,88 | 5,6 |
| 26 | Israel | 24 | 0,49% | 730 | 0,52% | ➡ | 534 | 0,56% | ➡ | 11,13 | 29,79 | 9,7 |
| 27 | Russian Federation | 24 | 0,49% | 541 | 0,38% | ➡ | 302 | 0,31% | ● | 7,04 | 41,87 | 7,4 |
| 28 | Ireland | 23 | 0,48% | 849 | 0,6% | ● | 563 | 0,59% | ● | 10,11 | 36,87 | 7,5 |
| 29 | Poland | 22 | 0,45% | 335 | 0,24% | ● | 247 | 0,26% | ● | 4,97 | 23,64 | 7,9 |
| 30 | Malaysia | 19 | 0,39% | 268 | 0,19% | ● ● | 180 | 0,19% | ● ● | 3,63 | 21,42 | 5,8 |
|  | **Totals/averages** | **4844** |  | **140810** |  |  | **96000** |  |  | **6,04** | **28,89** | **5,83** |

**Table 21.**
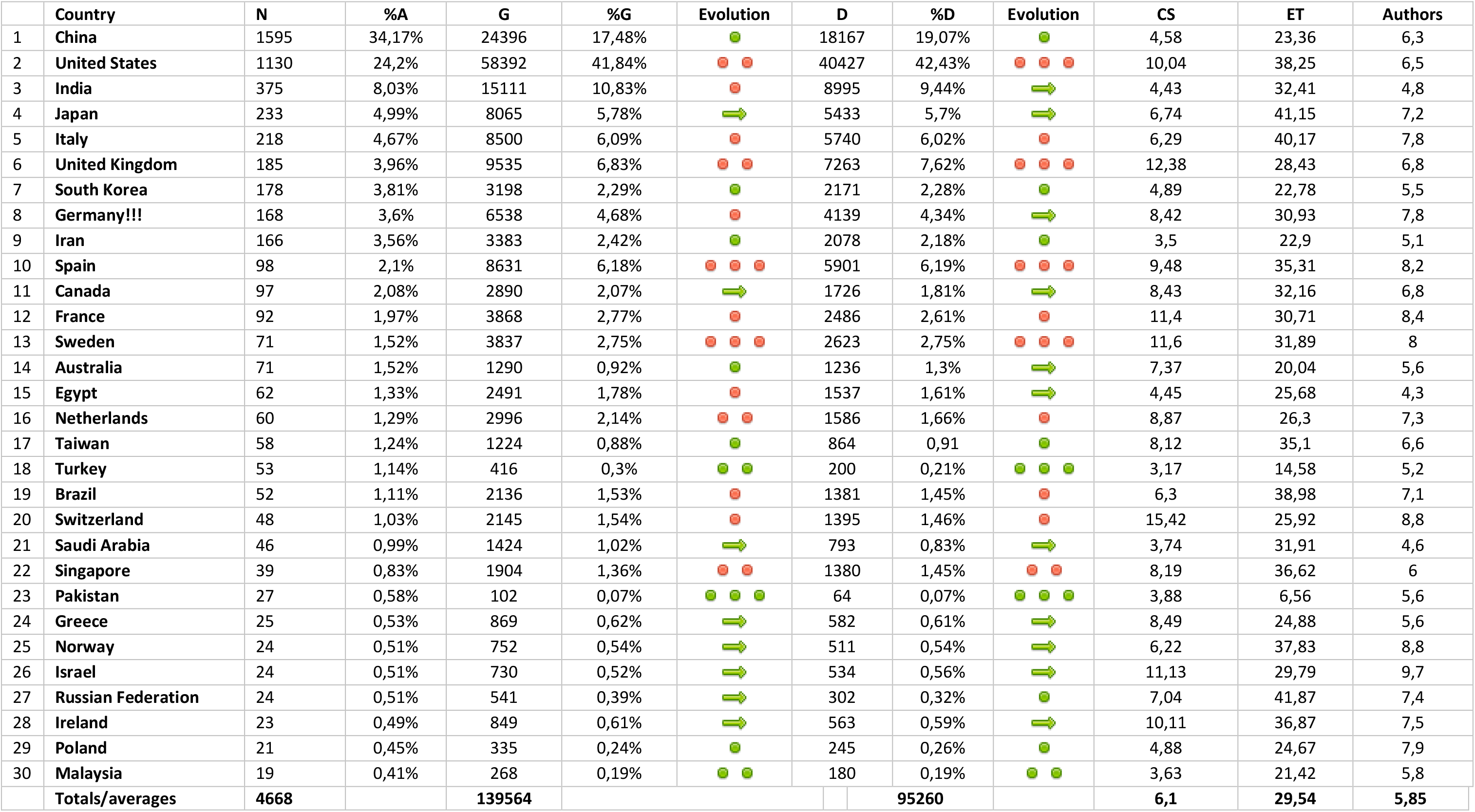
Retracted research impact for top 30 countries(all authors, without editorial retraction reason). 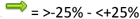; 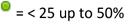; 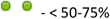; 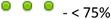; 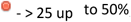; 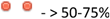; 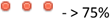; CS - CiteScore average(3933 records with available information) ; ET - Exposure Time average

The top 30 countries account for 94.7% of all withdrawn articles.

The data does not radically change when considering the country of origin of any author of the article. More consistent growth when compared to first author country numbers, reflecting probably a systemic penchant for international cooperation, is recorded in the United States, United Kingdom, Germany, and Canada(table 20).

### Retraction reasons

The withdrawal of a scientific article is a complex process in which several reasons can justify the retraction decision. In our series, we have found that 12,6% of the articles had multiple retraction reasons.

Mistakes and data inconsistencies represent 32,06%(1st place), the most affected countries being United States, United Kingdom, and Japan.

There is a substantial increase of scientific articles retracted because of image-related issues at a rate lower(22,46%) than the one estimated on a PubMed sample by Bucci [46], but still significant. Since the paper published by Rossner in 2004 [47], much progress has been made [48], but we can hypothesize that only the recent years’ advances in image processing and analysis have generated the adoption by publishers and journals of technologies able to discover image related QRP/QPP which were not easy to identify on a large scale a couple of years ago. The dynamic of retraction notes seem to support such a hypothesis (table 22). Countries with high image retractions percentages are the United States, Italy, and India.

**Table 22.** Evolution or retraction notes for images.

|  | Retraction note year |  |  |  |  |  |  |  |  |  |  |  |  | Total |
| --- | --- | --- | --- | --- | --- | --- | --- | --- | --- | --- | --- | --- | --- | --- |
|  | 2009 | 2010 | 2011 | 2012 | 2013 | 2014 | 2015 | 2016 | 2017 | 2018 | 2019 | 2020 | 2021 |  |
| Image retractions | 1 | 8 | 16 | 20 | 37 | 42 | 57 | 118 | 120 | 140 | 198 | 315 | 16 | 1088 |

Plagiarism and overlap continue to represent a problem with more than 25% of the retraction causes(plagiarism and overlap excluding images) and more than 30% when images overlap and plagiarism is added. India, Italy, and Iran have disproportionately large percentages(around 50%) of their retractions in this category.

The 5th place is the fraud, predominantly represented by fraudulent peer review(350 out of 393). Iran(32,7%), China(18,2%) and South Korea(16,9%) have important percentages of their retractions in this category. As the top years were 2015(118 retraction notes), 2016(75 retraction notes), and 2017(134 retraction notes), we have reasons to think that publishers and journals have fixed their vulnerabilities.

Ethics also represent an important reason for retraction, with 7,43% of the retracted articles. Interestingly enough, countries we did not expect have in this category substantial percentages of the volume of their withdrawn articles: Germany(20,9%), Japan(15,6%), South Korea(12%). Also, only one country in the top 10 is below 5%(Iran, 4,9%), and the number of retractions for ethical reasons is continuously increasing(Table 23). This fact leads us to consider the possibility of greater attention from publishers and journals to research ethics and/or publication ethics.

**Table 23.** Evolution of retraction notes for ethical reasons.

|  | Retraction note year |  |  |  |  |  |  |  |  |  |  |  |  | Total |
| --- | --- | --- | --- | --- | --- | --- | --- | --- | --- | --- | --- | --- | --- | --- |
|  | 2009 | 2010 | 2011 | 2012 | 2013 | 2014 | 2015 | 2016 | 2017 | 2018 | 2019 | 2020 | 2021 |  |
| Ethics | 1 | 5 | 21 | 18 | 20 | 27 | 23 | 25 | 33 | 41 | 56 | 88 | 2 | 360 |

Authorship is the retraction reason for 5,78% of the papers; editorial mistakes account for 3,73%, property & legal concerns 2,5%, and other reasons 1,3%. We must mention here that a relatively high number of papers(241, 5,1%) have no clear reason mentioned as the cause of retraction. Details about distribution by the top 10 countries are in Table 15.

The data do not change very much when the countries of all authors are taken into account(Table 16).

### The number of authors

Several published reports on the number of authors in the scientific literature [49–52] mention a continuous increase of average number. With an average of 5,83, a median of 5, and a mode of 4 authors per retracted article published between 2009 and 2020, we did not find an upward trend for the average number of authors. Single author papers account for 5,3% of the total, confirming the decreasing trend reported previously [50].

### Citations, journal impact, and exposure time

Citations and citations based indicators are still considered to reflect the impact and relevance of scientific work.

During the period in which the journal article is available both in search and on the journal’s website without any mention of withdrawal, it is considered valid by many researchers and can influence decisions related to research projects or, in the least pessimistic scenario, can be cited in the bibliography of an article.

Not all articles are cited, but there are enough situations in which the critical analysis of the content can be diminished in intensity by the journal’s prestige, author’s affiliations, or other factors. Therefore, to better highlight this multi-dimensional model, we extracted <u>citations</u> of withdrawn articles and the <u>journal’s impact indicator</u> (CiteScore) from the year of publication of the article (information available for the period 2011-2020). Also, the exposure time in months was calculated as described in the material and methods section for each article.

### Citations

For all 4844 articles, the total number of citations in Google Scholar is 140810(mean=29,07/median=11), and the total number of citations in Dimensions database is 96000(19,82/7).

We found a 68% coverage of Dimensions database when reported to Google Scholar, which was slightly higher than a recent report [39].

Most cited articles were the ones retracted for image reasons(average 44,6 on Google Scholar/32,9 on Dimensions) followed by mistakes(35,5/24,6) and plagiarism(27,8/16,1). Less cited were those retracted for editorial reasons(6,9/4,1), property and legal concerns(10,4/6,9) and authorship(11,4/7,2).

### Journal Impact (Cite Score)

There is old history and many controversies behind the indicators that were initially used to ease the purchasing decisions of academic libraries [53,54] or to assess the quality of scientific literature [55] and nowadays to give a measure of prestige for scientific journals [56]. Currently, there are two leading indicators for the impact of a journal: JIF(Journal Impact Factor, dominating the academic market since 1960) and CiteScore(launched in 2016) [57]. We extracted the journal CiteScore(CS) score(publicly available) for all retracted articles. CS information was available for 3933 articles published between 2011 and 2020.

Average CiteScore for 3933 articles is 6,03(5,81-6,26).

Articles with retractions reasons ,Other’, ,Mistakes/inconsistent data’ and ,lmages’ had the largest CS average value. In contrast, articles retracted for plagiarism, fraud, overlap, or authorship were published in lower CS journals (see table 12 for details).

### Exposure time

Exposure time was 28,89 months in average(median=19). Previous studies reported article lifespans between 26 months and 44 months [25,29–31,34,35,44] excepting the paper by Singh [37] which reports an 18 months lifespan for papers published between 2004 and 2013(12 months for 1695 papers published between 2009 and 2013). Our findings show a moderate decrease in the article lifespan for 2009-2020 compared to the previously reported data. For 2125 the papers published between 2009 and 2013, the lifespan is 41 months, a difference(compared to Singh) which late retractions for images/other reasons could explain.

Articles retracted due to image issues have a much longer average exposure time (49.21 months) than items withdrawn for any other reason (Table 12). This long exposure time, together with an average CS significantly higher than the average of the whole group and an average number of authors also higher than the average (the highest of all reasons for withdrawal), could explain why this group has the highest number of citations/article (44.6 in Google Scholar, 32,9 in Dimensions).

Articles withdrawn due to errors or inconsistency of data have a CS of 8.06 (highest) and an average number of authors of 6.4 (second in size) but an average exposure time of only 27 months. In this case, Google Scholar citations are 35.9 / article and 24.9 / article in Dimensions.

The other causes are significantly below average exposure time (less ,Ethics’ reason, with 31.7 months). All have fewer authors on average, and excepting the reason ,Others’, have a significantly lower CS.

### Impact of retracted research

We tried, in this article, to formulate a representation of the impact(citations received) that retracted research (all retraction reasons, without editorial reasons, country of the first author, countries of all authors) for the top 30 countries (tables 18,19,20,21).

### First author only, without editorial reasons(Table 19)

Our findings suggest that retracted scientific papers from 9 out of the first 30 countries have a higher impact (United States, India, United Kingdom, Spain, Egypt, Brazil, Netherlands, Singapore, Ireland) while retracted research from other countries has a lower than expected impact (Pakistan, China, Turkey, Malaysia, Russian Federation, South Korea, Iran, Australia, Saudi Arabia, Poland, and Greece).

### All authors, without editorial reasons (Table 21)

High impact for 13 countries: Spain, Sweden, United States, United Kingdom, Netherlands, Singapore, Switzerland, India, Italy, Germany, France, Egypt, Brazil;

Low impact: Pakistan, Turkey, Malaysia, China, South Korea, Iran, Australia, Taiwan, Poland.

Some outliers can bring some modifications. For example, Spain has an article with more than 4000 citations in Google Scholar. Australia has a highly cited article in Dimensions, while the same article has a small number of citations in Google Scholar. However, these do not modify the direction of the impact(Spain has a single red point instead of three, Australia loses one green point and passes to stable in only one of the four classifications).

Various factors may explain these differences in the impact of retracted research when considering the originating country: scientific tradition and strategies, funding of research, faster or easier access to better journals, scientific networking, better international cooperation, journal or institution prestige. When one or several of these factors concur in the direction of a negative impact of questionable research or writing practices, questions could be raised about the safety mechanisms of the scientific process.

The ecosystem of withdrawn articles is a complex one. The matrix in which they are framed reflects cultural differences, differences in development, and organization of the scientific research system in different countries that contribute to the shared heritage and advancement of biomedical knowledge. Our study is only a snapshot of a short period and its exploratory nature, and inherent weaknesses (lack of an unambiguous wording in some withdrawal notes, especially for ethical reasons; lack of content in over 5% of the withdrawal notes; online absence of some articles) determines us to proceed with caution in formulating conclusions.

## Conclusions

Mistakes and inconsistent data (including data fabrication) are the main retraction reason for articles published between 2009-2020 and indexed in PubMed.

Images and ethical issues are retraction reasons growing in recent years.

Plagiarism and overlap still represent a significant problem (> 30% of the total when images are included), especially in journals with a low impact (measured byCiteScore).

The number of fraudulent peer review cases shows the need to strengthen these processes, making them less vulnerable to circumvention attempts. As with plagiarism and overlap, the journals affected are those with a low CiteScore.

Major publishers and biomedical journals are involved in the retrospective verification (going back at least 12 years), resulting in a steady increase in withdrawals. The process does not appear to be slowing down.

Withdrawal of articles seems to be a technology-dependent process (image analysis and anti-plagiarism software). Increasing the accessibility of these technologies (price, usability, performance) can help combat QRP and QPP more effectively.

The number of citations of retracted articles shows a high impact of papers published by authors from certain countries, most of them developed. This impact suggests a need for improving the verification processes at the national and institutional levels, publishers, and biomedical journals.

The number of retracted articles per country does not always accurately reflect the scientific impact of QRP and QPP papers.

## Notes

### Competing Interest Statement

The authors have declared no competing interest.

### Summary of Updates

Revision 3(18.10.2021) - text reformatting, bibliography corrected and renumbered. Correction of data in paragraph 4, page 9: . Image reasons count 741(was 714) cases when image overlap and plagiarism are removed. Correction of data in paragraph 3, page 9: There were 229 instances of data fabrication, 217 in the "Mistakes/Inconsistent data" category, and 12(old value 10, not updated after quality check) in other categories. Replaced in abstract: "Mistakes and inconsistencies" with "Mistakes and data inconsistencies"

